# WildPose: A Long-Range 3D Wildlife Motion Capture System

**DOI:** 10.1101/2024.02.05.578861

**Authors:** Naoya Muramatsu, Sangyun Shin, Qianyi Deng, Andrew Markham, Amir Patel

## Abstract

Understanding and monitoring wildlife behavior is crucial in ecology and biomechanics, yet challenging due to the limitations of current methods. To address this issue, we introduce WildPose, a novel long-range motion capture system specifically tailored for free-ranging wildlife observation. This system combines an electronically controllable zoom-lens camera with a LiDAR to capture both 2D videos and 3D point cloud data, thereby allowing researchers to observe high-fidelity animal morphometrics, behavior and interactions in a completely remote manner. Field trials conducted in Kgalagadi Transfrontier Park have successfully demonstrated WildPose’s ability to quantify morphological features of different species, accurately track the 3D movements of a springbok herd over time, and observe the respiratory patterns of a distant lion. By facilitating non-intrusive, long-range 3D data collection, WildPose marks a significant complementary technique in ecological and biomechanical studies, offering new possibilities for conservation efforts and animal welfare, and enriching the prospects for interdisciplinary research.

## INTRODUCTION

Animal motion data play a pivotal role in multiple research disciplines, but current motion capture systems are largely confined to laboratory settings (Nagy et al., 2023). To achieve a holistic comprehension of natural animal behavior equivalent to the insights garnered within laboratory settings, the acquisition of high-resolution and precise 3D data becomes imperative. However, studying animals in their natural habitats introduces a unique set of challenges that continue to be actively addressed by researchers.

Existing methodologies for observing free-roaming animals employ various sensing techniques, each with their own strengths and limitations. GPS-IMU collars are effective for long-term tracking of animals over large areas and their inertial data is useful in maneuverability analyses (Wilson et al., 2013b; 2018). However, they require animal capture and sedation to fit, which can be costly, dangerous, and may influence natural behavior (Brooks et al., 2008; Coughlin and van Heezik, 2015). They also lack the ability to capture fine-grained body pose and surface details.

Camera traps have seen advancements using depth/stereo-camera data (Klasen and Steinhage, 2022) and vision-based post-processing (Pereira et al., 2020; Hernandez et al., 2024), but are restricted to high-traffic areas and provide limited 3D data. Static multi-camera field videography (Theriault et al., 2014) offers another approach, using fixed camera arrays to capture 3D data within the camera overlap zone, but lacks portability. UAVs equipped with cameras (Basu et al., 2019; Koger et al., 2023) are highly mobile but have limited flight time and may alter animal behavior due to noise when flying within 100 meters (Bennitt et al., 2019).

Zoom-lens photography is a common approach for convenient, high-resolution focal observations at long range. Using scene references, some morphometric data can be measured (Galbany et al., 2016; Brown and Wells, 2020; Cui et al., 2020; Breuer et al., 2007; Richardson et al., 2022; Rothman et al., 2008), but these methods fall short in capturing detailed 3D data like surface contours or body asymmetry. Telephoto lens stereo vision (de Margerie et al., 2015) enables 3D tracking of free-moving animals with a spatial uncertainty of < 0.1 m (root-mean-square) within 100 m of the observer, sufficient for gross motion tracking but lacking the spatial resolution for detailed whole-body kinematics.

In summary, existing approaches are subject to trade-offs between tracking range, duration, 3D data resolution, and morphometric detail (Table S1). Each method has its strengths in certain aspects of wildlife observation, but no single approach currently provides a comprehensive solution for high-resolution, long-range 3D motion capture in natural habitats.

In light of these limitations, we introduce WildPose, a novel motion capture system specifically tailored for 3D wildlife observation in natural environments (Fig. 1A). Although a stereo camera system is suitable for animal observation in the field, the position error increases in proportion to the square of the distance from the target object (de Margerie et al., 2015), so our system used a light detection and ranging (LiDAR) sensor which can potentially measure distance with a stable, low error (e.g. below 0.05 m from 5 to 180 m) (Lambert et al., 2020).

**Fig. 1.**
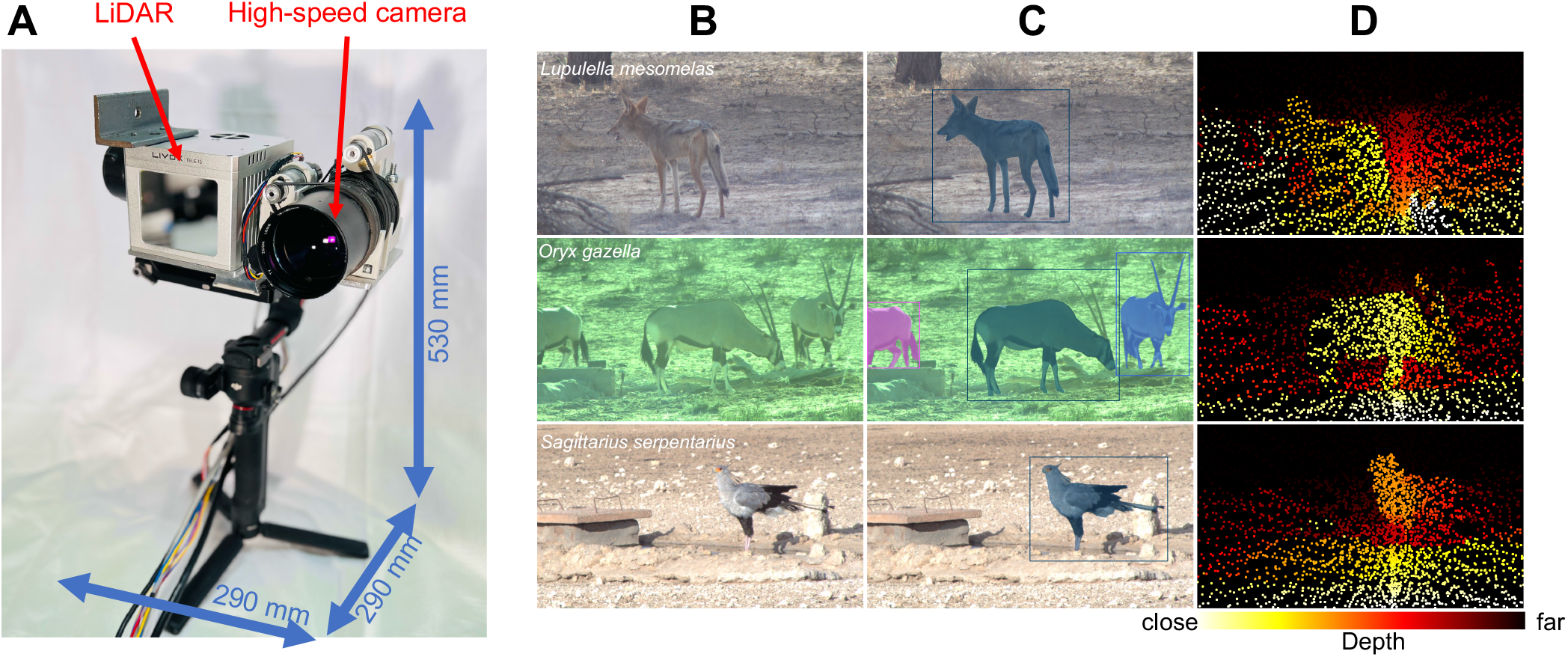
The WildPose long-range 3D motion capture system. (A) The main hardware components of the WildPose system, which was mounted on a car door in field work. (B–D) Example frames with jackal, gemsbok, secretary bird, and red hartebeest. Given a zoom-lens camera image (B), the image foundation model (Kirillov et al., 2023) outputs segmentation masks of the wild animals with 2D bounding box annotations (C). The corresponding point cloud projected onto the image plane, with colors representing the depth from the image sensor (D). Note that the LiDAR provides full 3D information (x, y, z) derived from LiDAR angle and depth measurements.

**Fig. 2.**
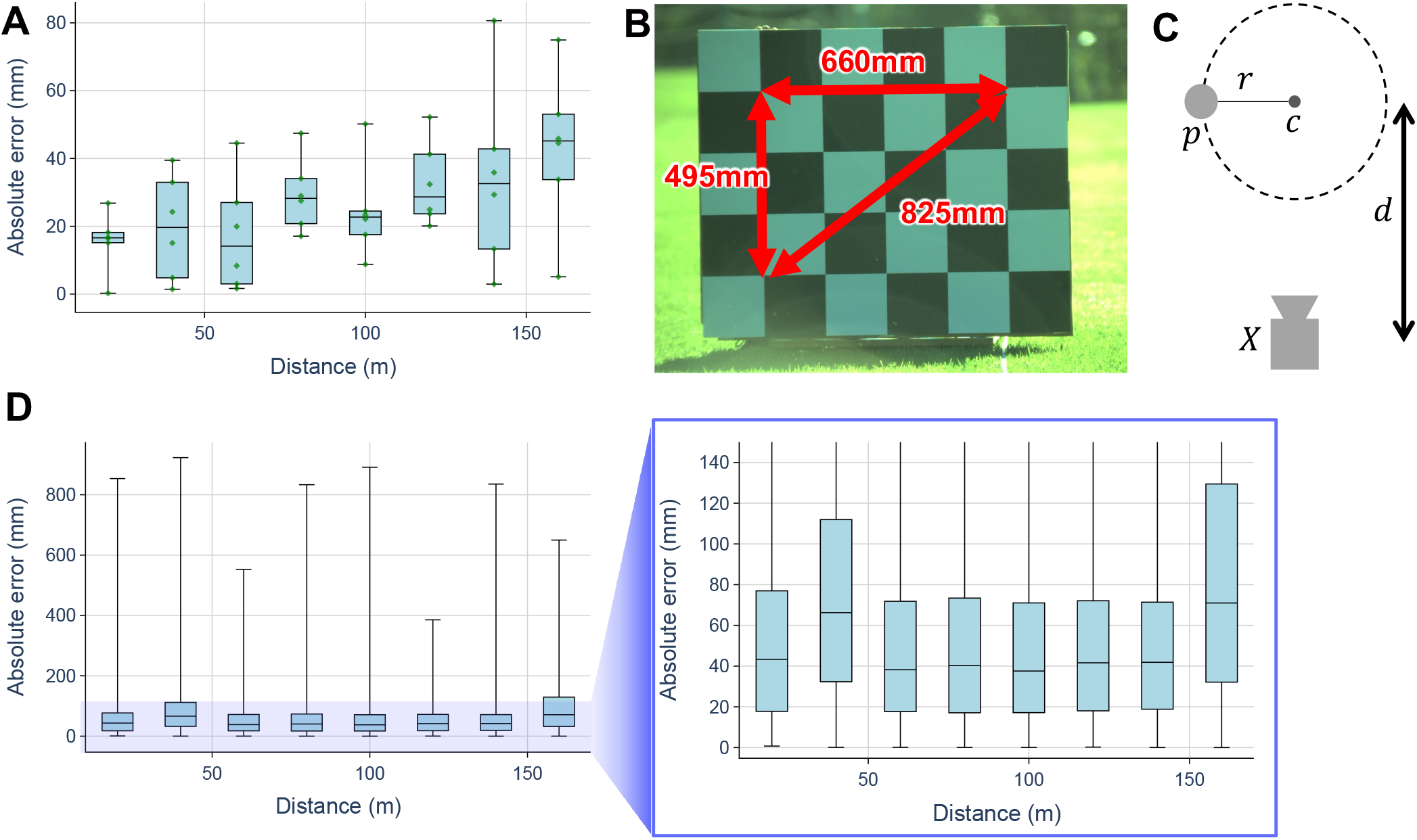
Validation and capabilities of the WildPose system. (A) Absolute error of measured lengths on a checkerboard at various distances. X-axis: distance to board (m); Y-axis: absolute error (mm). Box plots show error distributions: median (central line), 25th-75th percentiles (box), full range excluding outliers (whiskers). Green dots represent individual measurements. (B) Reference lengths on the checkerboard used for measurements.(C)Experimental setup for tracking a moving subject. Subject *p* walked in circles of radii *r* around a pole *c. d* is the distance between the system *X* and pole *c*. (D) Absolute error in tracking a moving subject at various distances. Plot structure similar to (A), showing error distributions for different circular paths. Left panel: full range of errors; Right panel: zoomed-in view of smaller errors.

The WildPose system meets three critical criteria and provides additional functionalities:

(i) 3D Data Acquisition: Vital for modern biomechanics research (Nourizonoz et al., 2020; Muramatsu et al., 2022; Lobato-Rios et al., 2022; Peng et al., 2020).
(ii) Long-range Capability: Designed to minimize behavioral influence, allowing data capture from considerable distances.
(iii) Portability: Easy to transport and set up in field works.

Unlike existing solutions, WildPose features a compact, mobile design optimized for unobtrusive data collection in challenging terrains. As shown in Table S2, existing popular, portable 3D scanners have difficulty observing animals at a distance, but WildPose can acquire detailed 3D information from much longer distances.

We validated the system’s capabilities through field tests, capturing a diverse range of animals and generating colored point clouds.

This paper further explores the implications of WildPose for wildlife research and identifies potential directions for future enhancements. One notable application is the precise measurement of animal morphometrics, achievable with a precision of less than 50mm, across a broad range of species — from giraffe to eagles. Precise individual tracking of animals, e.g., within herds, is another application of WildPose. In a case study involving a springbok herd, WildPose captured intricate behaviors, such as sparring, from a distance of approximately 150 meters. Leveraging the image foundation model, Segment Anything (Kirillov et al., 2023; Ravi et al., 2024), our system employs high-resolution video data for automated individual animal segmentation. Integration with the corresponding 3D points enables precise tracking of individual movements with a typical 3D positional error of ± 42mm at 10 H_z_. This error includes both precision (random error) and accuracy (systematic error), and is considerably lower than the typical total error of GPS collars, which can range from 1.8 to 5 meters depending on the environment and GPS receiver quality (Wilson et al., 2013a; Wilmers et al., 2015; Foley and Sillero-Zubiri, 2020). Moreover, as a novel example, we show how WildPose can be used to capture subtle body movements, such as lion’s chest motion during sleep. Through these example applications, we demonstrate that WildPose constitutes a significant step forward in animal behavior research, offering a unique and effective tool for high-quality 3D data acquisition in natural habitats.

## MATERIALS AND METHODS

### Implementation of the WildPose system

WildPose integrates LiDAR technology with a zoom-lens high-speed camera, both enhanced by an active gimbal for remote wildlife observation in natural settings (Fig. 1A). The long-range sensing capability minimizes human impact on animal behavior, providing a more authentic observational context without disturbance. The use of LiDAR point cloud ensures the potential acquisition of metric data which is indispensable for analyzing inter-individual interactions and making quantifiable, repeatable morphological measurements. Additionally, a rugged design coupled with an active gimbal enables remote control of pan and tilt, ensuring operator safety during fieldwork. We substantiated the system’s capabilities through field trials in Kgalagadi Transfrontier Park (South Africa), capturing a wide spectrum of animal data.

LiDAR serves as a high-quality depth sensor. Due to its repetitive scanning pattern of mechanical spinning LiDAR, e.g., Velodyne (https://velodynelidar.com/), the reflected point cloud tends to be sparse for the interest area. To address the sparsity concerns associated with traditional LiDAR systems, we selected a prism-scanning LiDAR from Livox (https://www.livoxtech.com/), which provides dense point clouds with a narrow field of view (Liu et al., 2022). In particular, the device features a unique scanning mechanism that produces non-repetitive point cloud patterns, improving the richness of the data captured compared with traditional line scanning LiDARs. For this study, we used the TELE-15 model, which has the narrowest field of view (14.5^°^ × 16.2^°^) in the product series and a high density of points (480, 000 points/s). It has the potential to capture 3D points in metric system with the depth precision of less than 20mm under optimal conditions (https://www.livoxtech.com/tele-15/downloads). However, real-world performance may vary from these specifications, as demonstrated in previous work (Lambert et al., 2020). Therefore, we tested the precision of the entire system under conditions that approximated real applications, as detailed in the “Static measurement precision” and “Object tracking precision” sections.

The zoom lens (effective focal length: 28.35–283.5mm) camera complements the LiDAR by providing high-resolution visual data, which combines two crucial features: variable focal length (zoom capability) and high magnification at long focal lengths (telephoto capability). It allows for flexible data collection at various distances and long-range observations. Motors and encoders were affixed to the lens for remote zoom control, allowing for rapid adjustments to capture both wide-angle contextual shots and telephoto close-ups of distant subjects. This flexibility is crucial for adapting to dynamic wildlife scenarios. Representative frames of the camera and LiDAR illustrating this are depicted in Fig. 1B–D.

To orchestrate the multitude of sensors and actuators in WildPose, we built the software architecture on ROS2 (Macenski et al., 2022). However, we substituted the default ROS2 middleware, Real-Time Publish Subscribe Data Distribution Service (RTPS DDS), with eCAL RMW (https://github.com/eclipse-ecal/rmw_ecal) to facilitate quicker inter-process communication. This decision was motivated by the inefficiency of RTPS DDS in long-duration data recording, which led to frame loss (see the “eCAL RMW” section). On the other hand, eCAL RMW prioritizes performance and simplicity, even though this comes at the expense of increased storage requirements. This is the key software design to record standard high definition video (resolution of 720 × 1280) with 170 FPS and LiDAR points with 10 FPS. In summary, the WildPose system accurately records the animal’s 3D data even at a long distance, and provides a high-resolution close-up view of the animal’s behavior.

#### Hardware setup

As shown in Fig. S2, WildPose consists of stabilized multisensor module mounted on a car door during the field work and the control module including the display, gamepad controller, mini-computer and electrical circuits.

The multi-sensor module has three main components: a LiDAR, motorized zoom-lens camera and active gimbal. As the LiDAR, we used Livox TELE-15 (recorded at 10 FPS) for the narrow field-of-view (14.5^°^ × 16.2^°^) is suitable for combining with zoom-lens camera. A 170 FPS camera (MQ022CG-CM, XIMEA) was used for capturing the video data, whose sensor active area is 11.27 × 6mm. The cameras were equipped with zoom lens (Super16 Zoom Camera Lens, 16–160mm FL, NAVITAR), which is controlled by three encoder-attached DC motors (9.7:1 Metal Gearmotor 25D × 63Lmm HP 12V with 48 CPR Encoder, Pololu) through pulleys. The theoretical field of view of the camera is from 43.2^°^ × 22.7^°^ to 4.5^°^ × 2.3^°^. The motorized zoom lens system allows for remote control and electronic recording of the lens status from a safe location, such as inside a car. However, the motors and pulleys introduce lag, play, and hysteresis in the control system. Therefore, the pulley positions were reset for each session during field work, and the system’s lens status was used as a reference for camera parameter tuning rather than an accurate measurement.

To automatically stabilize the sensor system on a car, we used DJI RS3 Pro, which holds the LiDAR and motorized zoom-lens camera. The focus wheel attached on the gimbal provides the CAN interface to the main computer, allowing for remote control of the pan and tilt.

The control module consists of two computers. As the main computer recording all the sensor data, controlling the active gimbal and display the visual status of the system, we used NVIDIA Jetson AGX Orin Development Kit with 1TB V-NAND SSD (970 EVO Plus NVMe M.2 2280, Samsung). Teensy 3.2 microcomputer board was used for controlling a motor driver of the zoom lens, correspond to commands from the main computer. The operator can control the zoom-lens parameters (focus, zoom level and aperture), gimbal posture (yaw and pitch), recording states using by a gamepad controller (F710 Wireless Gamepad, Logitech).

#### Software setup

The previously mentioned sensors are seamlessly integrated into a compact and robust platform. This platform operates on an Nvidia Jetson AGX Orin ensuring efficient capturing and synchronization of sensor data with high throughput. The WildPose software, which is built upon ROS2 Foxy, utilized a combination of C++ and Python. For 3D visualization of data points, we harnessed the capabilities of Open3D (www.open3d.org). The image sensor system was managed using XiAPI libraries for C++ (www.ximea.com/support/wiki/apis/XiAPI). Additionally, a Teensy 3.2 board, programmed with the Arduino IDE, was implemented to manage zoom-lens motor rotations and record encoder steps.

#### eCAL RMW

Capturing high-speed HD video alongside 240,000 points per second LiDAR data with ROS2 poses a significant challenge in our software system. The standard ROS2 RTPS DDS often grapples with being resource-intensive and exhibits bandwidth limitations and latency issues (Kronauer et al., 2021). To overcome this hurdle, we employed eCAL RMW, an efficient ROS2 middleware developed from the open-source eCAL framework (https://github.com/eclipse-ecal/rmw_ecal). This middleware enables streamlined communication among ROS2 components utilizing the eCAL framework. Contrasting the ROS2 RTPS DDS, eCAL RMW underscores simplicity and peak performance. It diverges from adherence to any DDS standard, instead concentrating on fostering efficient data communication, reducing storage and bandwidth requirements, and ensuring low-latency data transfe (https://github.com/eclipse-ecal/ecal).

For data storage, we opted for the HDF5 file format, due to its enhanced accessibility relative to ROS2 Bags. Serialized data from each sensor is directly streamed into HDF5 files via an eCAL subscriber, allocating one subscriber per sensor topic. Subsequently, this data is deserialized and translated into human-readable formats, such as images and PCD, providing a comprehensive representation of the dataset.

#### Synchronization

Synchronization is a critical component for constructing an integrated multi-modal animal dataset. All sensor data in our system are timestamped at every frame. These timestamps are instrumental in identifying the nearest frame across sensors to construct a cohesive multi-modal frame. Notably, the camera video operates at 170 FPS, thereby limiting the maximum error margin with LiDAR data to 3 milliseconds.

### LiDAR-camera calibration and point cloud colorization

To the best of our knowledge, no established automatic method exists for calibrating a narrow field-of-view (FOV) camera with LiDAR point cloud data. Given the dynamic nature of camera parameters during field data collection, we employed a manual calibration approach for intrinsic camera parameters (Hartley and Zisserman, 2004).

We developed and utilized an in-house visualization tool (https://github.com/DenDen047/WildPose_v1.1/tree/main/calibrator) that projects the 3D point cloud from LiDAR onto corresponding image frames. This process involves estimating the intrinsic parameters of a pin-hole camera model (focal length and principal point offset) which cannot be precisely measured but rather estimated to allow for accurate alignment between LiDAR and camera data. Operators fine-tuned the parameters by leveraging laser reflectivity and depth cues from the camera images. The extrinsic parameters, defining the spatial relationship between the LiDAR and camera, were derived from the hardware CAD data, ensuring consistency across different setups.

To address the challenge of changing intrinsic parameters in the field, we maintained consistent lens settings for as long as possible during each data collection session. Calibration was performed for each scene with different lens settings, ensuring accurate alignment between LiDAR and camera data throughout the varying field conditions.

The colored point clouds were created by merging the calibrated camera and LiDAR data using the estimated camera parameters from the manual calibration process. Note that while the operator can confirm no-colored point cloud in the field during data collection, the generation of colored point clouds was performed offline.

### Animal extraction pipeline

The accurate extraction of animal bodies from a scene, complete with precise 3D coordinates, forms the cornerstone for leveraging both camera and LiDAR data in the WildPose system. We address this segmentation challenge by combining machine learning models with the calibrated 3D sensors capable of accurately detecting animals at a distance.

In the initial phase, 2D bounding boxes are annotated on the first frames of a monocular RGB image. These bounding boxes are subsequently propagated to successive frames using an image-based box tracking method (Redmon and Farhadi, 2018), facilitated by DeepLabel (Veitch-Michaelis, 2021). This methodology substantially expedites the annotation process, as it obviates the need for extensive manual labeling beyond the first frame. However, 2D bounding boxes offer only a coarse-grained representation, encapsulating merely the center, width, and height of the object while treating it as a rigid body. Additionally, these boxes often encompass background areas, thereby introducing extraneous data. To refine this representation, we employ Segment Anything (Kirillov et al., 2023), a foundational model for image segmentation, as the next stage in our pipeline. Using the 2D bounding boxes generated in the first phase as input, Segment Anything produces finely-tuned segmentation masks for the predominant object within each box, as illustrated in Fig. 1C.

The final step in the process involves back-projecting these foreground pixels from the segmentation masks to 3D coordinates in the camera’s frame of reference. To establish the correspondence between points in the camera coordinates and pixels in the image plane, we first transform points from the LiDAR coordinate system to the camera coordinate system using known extrinsic parameters. These transformed points are subsequently projected onto pixel coordinates through perspective projection, employing estimated intrinsic and extrinsic parameters as shown in the “LiDAR-camera calibration and point cloud colorization” section. Following this projection, we retain only those pixels that have a direct mapping to the 3D space (Fig. 3A and Fig. S1).

**Fig. 3.**
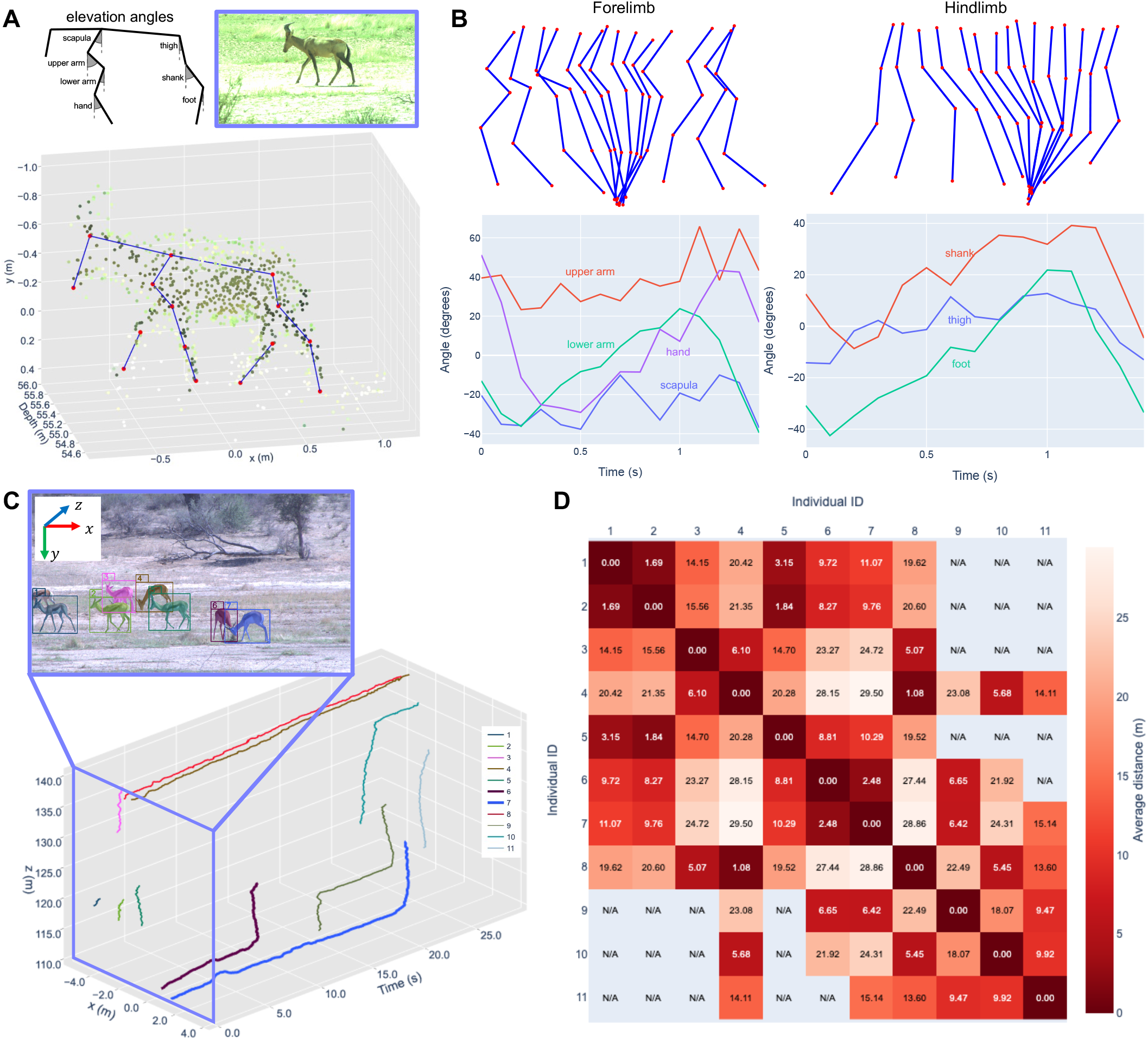
Measurement of 3D animal motion. (A) A red hartebeest walking scene captured over 1.4 seconds. Top-left: Segment definitions for elevation angles. Top-right: A sample camera frame. Bottom: Corresponding 3D point cloud of the sample frame reconstructed with tracked limb segments in the camera coordinate system. (B) Limb segment elevation angles. Top: Angular movements along the sagittal plane for both forelimb and hindlimb, with red dots indicating keypoint positions and blue lines connecting them. Bottom: Plots of each angle over time, with positive direction defined as anticlockwise. (C) 3D trajectories of the individual springboks in a herd, where the z-axis represents the depth information from the camera, and the camera frame corresponding with the purple plane. After the springboks (ID 6 and 7) were sparring in the first time (*t* = 0 s), ID 6 was the first to leave (*t* = 14 s). (C)Heat map of the average distances between each pair of individual springboks, where warmer colors indicate shorter distances and cooler colors indicate longer distances. “N/A” means that both individuals were not present in the frame simultaneously.

Through this multi-step process, we successfully obtain accurate 3D segmented masks of animals. This segmentation methodology serves as a foundational approach for various downstream analyses, which we elaborate upon in the subsequent sections.

### Data collection in Kgalagadi Transfrontier Park

In December 2022, we embarked on a wildlife recording expedition in Kgalagadi Transfrontier Park, South Africa, using the WildPose system mounted on a vehicle door. Over the course of 13 days, we conducted regular surveys along fixed routes within the road network surrounding the Two Rivers and Nossob camp sites. Whenever land animals or large birds were encountered, we captured their movements for duration ranging from 30 to 900 seconds. To keep image brightness at a fixed frame rate (170 fps) and shutter speed (1000 ms), we adjusted the zoom lens aperture according to the varying lighting conditions.

### Measurement of morphometrics

To measure of morphometrical data of animals, we identified and defined four critical keypoints for this analysis: `nose`, `r_eye` (right eye), `neck`, and `hip` as shown in Fig. S1. Here, we measured two animals, martial eagle and giraffe, to prove the robustness of the procedure for body size.

For the martial eagle, the `nose` keypoint corresponds to the tip of its beak. The `r_eye` keypoint is situated at the right eye. The `neck` keypoint is located at the base of the neck. Finally, the `hip` keypoint is identified as the base of the tail in the giraffe and the tip of the tail in the martial eagle. When possible, we measured the following features: overall height (ground to parietal), head length (`nose` to `r_eye`), neck length (`r_eye` to `neck`) and body length (`neck` to `hip`). For a more detailed locomotion analysis, we expanded our approach to a walking red hartebeest. In this case, we defined a comprehensive set of 21 keypoints (Fig. 3A).

These points were initially identified manually on 2D video frames, with their corresponding 3D positions marked on the LiDAR frame. While keypoints other than `nose` were designated on both the right and left sides, occlusions often obscured one side in most frames, highlighting a common challenge in field-based biomechanical studies.

### Statistics and reproducibility

No data were excluded from the analyses. All values reported as mean ± SD unless otherwise stated. Plot and analyses were performed in Python 3.8.19 (with the aid of numpy 1.24.4 and matplotlib 3.5.3).

## RESULTS AND DISCUSSION

### Static measurement precision

To evaluate the calibration accuracy of the WildPose system, we measured known lengths on a standard 5 × 6 checkerboard pattern with dimensions of 825 × 990mm and no margins (Fig. 2B). The checkerboard provides a flat, rigid surface with precise dimensions for assessing the system’s measurement capabilities.

After calibrating the LiDAR and camera as described in the “LiDAR-camera calibration and point cloud colorization” section, we collected data with the checkerboard positioned at 8 distances from the WildPose system: 20, 40, 60, 80, 100, 120, 140, and 160 meters. At each distance, the checkerboard was oriented at three angles relative to the line of sight of the WildPose system: perpendicular, − 45°, and + 45°. This setup enabled us to evaluate the system’s performance under varying orientations and distances.

For each session, we identified three pairs of checker pattern points in the camera image frame, representing three different length measurements across the checkerboard surface: 660, 495, and 825mm (Fig. 2B). We then located the corresponding 3D points within the LiDAR point cloud data and computed the Euclidean distance between them in 3D.

Fig. 2A displays the absolute error of the measured lengths compared to the true lengths for each point pair, plotted against the checkerboard distance from the WildPose system. The results demonstrate that the system can measure lengths with a median absolute error of 45.2mm at 160 meters. While there is a slight upward trend in absolute error with distance, the difference is minimal relative to the target distance. These findings align with previous literature (Lambert et al., 2020), suggesting that measurement accuracy remains relatively stable across the tested range.

### Object tracking precision

To assess the WildPose system’s positional accuracy for moving objects, we measured a human subject walking in circles with radii *r* of 1, 2, and 3 meters. The subject walked around a pole, connected by a string to maintain a constant radius (Fig. 2C). We collected data with the pole positioned at the same 8 distances used in the previous section, with the subject completing two revolutions at each setting.

For each scene, we manually selected a human segment in the first image frame, and SAM2 automatically generated segmentation masks for subsequent frames (Ravi et al., 2024). After visual verification of the masks, we calculated the 3D position of the person in each frame by projecting the center of the 2D mask onto the 3D point cloud. To account for potential inaccuracies in the string length and to handle outliers, we fitted a circle to the extracted trajectory using a least-squares method with RANSAC algorithm. We then calculated the absolute error between the fitted circle and each point. Fig. 2D shows the absolute error for each of the three circle settings, plotted against the pole distance from the WildPose system.

The results demonstrate a median absolute error of less than 71mm, even at distances up to 160 meters. Similar to the static measurements in the previous section, the positional accuracy remains relatively consistent across the tested distances. The slightly larger errors observed in this experiment compared to the checkerboard measurements can be attributed to factors such as the subject’s inability to maintain a perfect circular path and the thickness of the human body. Notably, the median error values in Fig. 2D (dynamic measurements) are not significantly different from the results in Fig. 2A (static measurements), potentially indicating consistent performance across both static and dynamic scenarios. Notably, the median error values in Fig. 2D (dynamic measurements) are comparable to those in Fig. 2A (static measurements), suggesting that the system may maintain similar levels of performance across both static and dynamic scenarios.

### Animal morphology & locomotion

WildPose’s first application aims to radically enhance the remote measurement capabilities for wildlife morphological features, which are invaluable for a wide array of biological investigations. Typical methods to measure the key characteristics of living animal body is to use invasively tranquilized animals, which is dangerous and risky for the animals and investigators.

Conventional long-range morphological measurements in the wild primarily utilize digital photogrammetry, broadly classified into two approaches. The first involves a distance meter, where measurements require precise distance determination between the camera and the animal, typically using a laser range finder (Brown and Wells, 2020; Cui et al., 2020; Breuer et al., 2007). The second method employs parallel lasers, projecting equidistant laser points onto the target for calibration (Richardson et al., 2022; Rothman et al., 2008). Both techniques, however, oversimplify by treating the animal’s body as a vertical flat plane, affecting accuracy.

WildPose leverages LiDAR technology to capture metrically accurate point cloud data of animals, with point density increasing if the animal remains stationary. To prove the validation for morphometrics measurement, we measured the 3D lengths of body parts of a giraffe and martial eagle from 2D keypoints. The measured lengths are catadepth errors are less than loged in Table 1 and the textured 3D point cloud data are shown in Fig. S1. The system’s long-range performance has allowed even small animals such as the martial eagle (Fig. 1) to be measured from a distance of 18.5 meters with an precision (standard deviation around mean measured position in repeated measures) of less than 11.2mm.

**Table 1.**
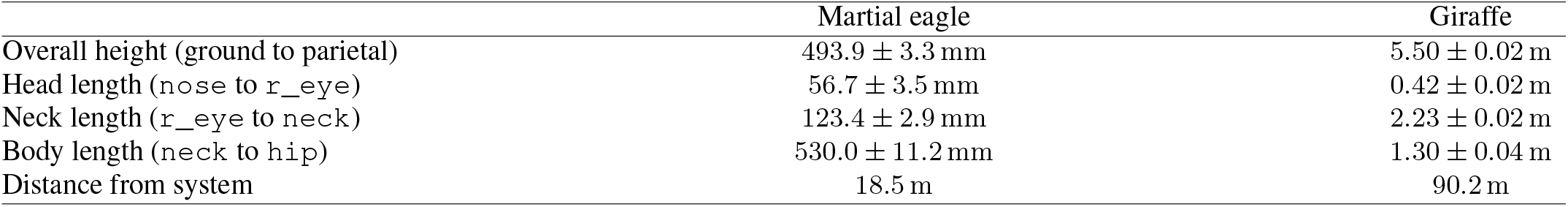
Morphometrics of wild animals. Each item indicates the measured mean and standard deviation of 17 measurements of the distance between key points. The definitions of keypoint are detailed in Fig. S1.

|  | Martial eagle | Giraffe |
| --- | --- | --- |
| Overall height (ground to parietal) | 493.9 ± 3.3 mm | 5.50 ± 0.02 m |
| Head length (nose to r_eye) | 56.7 ± 3.5 mm | 0.42 ± 0.02 m |
| Neck length (r_eye to neck) | 123.4 ± 2.9 mm | 2.23 ± 0.02 m |
| Body length (neck to hip) | 530.0 ± 11.2 mm | 1.30 ± 0.04 m |
| Distance from system | 18.5 m | 90.2 m |

While we did not have ground truth data to directly assess the accuracy (systematic error) of the measurements, LiDAR point clouds are generally more accurate than other methods such as zoom-lens stereo vision. Typical LiDAR depth errors are less than 0.25m mean residual error and 0.25m horizontal RMSE (Lambert et al., 2020). In contrast, the theoretical distance error for zoom-lens stereo vision with the same lens specifications is 900mm at 100 m distance (de Margerie et al., 2015). This suggests that depth inaccuracies are likely a less significant concern for the LiDAR-based WildPose system compared to alternative approaches. In summary, the precision (random error) of the WildPose measurements was quantified as less than 11.2mm standard deviation in an example case. While the absolute accuracy could not be directly measured, typical accuracy values reported for LiDAR systems indicate that systematic depth errors are likely small relative to the measurement scale from Fig. 2A.

Furthermore, the challenge of 3D pose estimation is critical, with various methodologies being proposed in numerous studies (Nath et al., 2019; Bala et al., 2020; Karashchuk et al., 2021). Our system, which merges 3D point cloud data with camera frames, advances the estimation of dynamic wildlife poses in three dimensions by utilizing scene data alongside 2D keypoint labels.

Fig. 3A and B demonstrate the system’s capability to capture and analyze a red hartebeest’s straight walk, showcasing both 2D and 3D representations of joint positions over time. We manually tracked keypoints in the camera frames and projected them into the 3D point cloud. The keypoint definitions on the legs follow those used by (Catavitello et al., 2018). Despite the animal walking at an angle of approximately 30° from the camera sensor and the short 1.4-second recording duration, we successfully reconstructed the leg segment motions from 3D point cloud data along the walking direction. Fig. 3B shows the 2D leg segment motions projected onto the sagittal plane. The metric data from the LiDAR point cloud allowed us to measure key locomotion parameters: walk speed (0.844 m/s), stride length (1.106m), leg length (0.82m), and Froude number (0.089). These values align with previous findings (Alexander and Jayes, 1983; Catavitello et al., 2018).

The angular changes of each limb segment over time are illustrated in Fig. 3B, demonstrating the system’s potential for detailed kinematic analysis. The plots are likely to show phase shifts between the upper arm, lower arm, and hand in the forelimb, as well as between the foot and shank in the hindlimb. These patterns are consistent with those reported by (Catavitello et al., 2018), although the short duration of our scene introduces some noise to the data. The elevation angle plots provide insights into the gait characteristics:

- Forelimb dynamics: The upper arm shows a larger range of motion compared to the lower arm and hand, indicating its primary role in propulsion.
- Hindlimb patterns: The thigh and shank exhibit similar ranges of motion, while the foot shows more pronounced angular changes, reflecting its importance in push-off and energy transfer.
- Temporal coordination: Phase differences between segments are visible, particularly between the upper and lower limb segments, illustrating the complex coordination required for efficient locomotion.

These detailed 3D joint angle analysis demonstrate the system’s potential for comprehensive biomechanical studies of wildlife in their natural habitats. By combining spatial and temporal data, our method enables researchers to quantify subtle aspects of animal locomotion that were previously challenging to measure in field conditions. However, the current setup has limitations. The single viewpoint results in occlusions, preventing the marking of some keypoints in the 3D visualization.

### Tracking individual animals in 3D

The ability to track the movements of wild animals in three dimensions is critical for addressing a diverse array of questions in animal ecology, behavior, and cognition. Long-term tracking, spanning days or even longer, provides ecologists with insights into large-scale spatial behaviors such as home ranges, migration patterns, and dispersal mechanisms. This data is typically provided by GPS-IMU collar modules. On the other hand, short-term and high-resolution tracking illuminates localized pathways of many animals during specific activity phases (Wilson et al., 2018). This granularity of data is invaluable for investigating an animal’s exploratory strategy, orientation skills, and biomechanical interactions with its environment.

WildPose provides animal location information that is fundamentally different from GPS-IMU modules by integrating LiDAR with zoom-lens cameras and post-processing techniques described in the “Animal extraction pipeline” section. It is particularly adept at tracking localized, large-scale animal behavior, such as the interactions in a herd. This capability is practically challenging to achieve using traditional collar-based methods.

Fig. 3C demonstrates this capability by tracking the centers of segmented 3D animal point clouds along with their headings in bird-eye-view (top-down) over time. The result achieves an average precision of 42mm, as measured by stationary individuals (ID 4 and 8 from 3.3 to 9.9 seconds in Fig. 3C), with individual positional errors summarized in Table S3. To showcase the ecological data captured and analyzed by our system, we have developed a heatmap visualization (Fig. 3D and Movie 1) that displays the average distances between each pair of individuals over time, effectively demonstrating interactions between individual springboks and providing valuable insights into their group dynamics and spatial relationships.

The WildPose system’s advantages are evident in scenarios where traditional approaches may fall short. For instance, while most springboks move in the negative x-axis direction, two specific individuals (ID 6 and 7 in Fig. 3B) initially spar and then diverge at different times. The heatmap visualization (Movie 1) clearly shows these interactions and separations over time. Capturing such nuanced behaviors with GPS-IMU modules (Grünewälder et al., 2012) would require sedating and instrumenting several animals.

The combination of precise 3D tracking and inter-individual distance visualization demonstrates WildPose’s unique capabilities in capturing and analyzing complex group dynamics and individual behaviors in wild animals. This approach opens up new possibilities for studying fine-scale interactions and movement patterns in natural settings.

While these capabilities offer significant advantages, it is important to acknowledge key limitations of our technique. Due to the narrow FOV as shown in Table S2, the volume of interest within which animals can be tracked continuously is relatively small. This constraint translates into short tracking durations, which may limit the ability to observe long-term behavioral patterns or movements over extended periods. These limitations particularly affect studies requiring continuous observation of wideranging behaviors or long-term movement patterns. Additionally, the photographer-style data collection method can be challenging for continuous recording due to obstacles like bushes in natural environments, potentially leading to data gaps in complex terrains. Despite these constraints, WildPose provides valuable insights into fine-scale animal behavior and group dynamics that complement traditional long-term tracking methods.

### Fine scale deformation monitoring

The third example application of WildPose serves as an innovative instrument for the nuanced study of breathing patterns in wildlife, an essential indicator of animal well-being tightly correlated with energetic expenditure across taxa (Ramanathan, 1964). By leveraging LiDAR technology, WildPose enables a pioneering method for remotely tracking detailed breathing dynamics. Building on the preliminary research that explored non-invasive remote vital data collection using thermal imagers (Rzucidlo et al., 2023), WildPose expands these capabilities, offering unprecedented granularity and accuracy in monitoring both the rate and amplitude of respiration in wildlife by measuring the expansion distance of the chest. This is notably demonstrated in the system’s ability to capture subtle thoracic motions in a resting lion from 31 meters away, a task traditionally constrained by the requirement for close proximity.

To measure the thorax height of a lion resting on its side, we first estimated the ground plane from the point cloud data. As the lion was situated on a flat road, we aggregated point cloud frames across the scene and employed the RANSAC algorithm to identify the largest plane as the ground (Torr and Zisserman, 2000). The chest height of the lion was then defined as the average height of 3D points in a frame relative to this identified ground plane. To isolate the lion’s breathing frequency amid data noise, a band-pass filter of 1–2 Hz was applied based on (Rzucidlo et al., 2023).

Fig. 4A illustrate the result of this process, as the derived size changes in the lion’s body align closely with manual observations. For validation, we manually annotated inhalation and exhalation phases. These annotations matched 96% of the local minima and maxima in the extracted respiration pattern. The filtered breathing pattern not only corroborated these labels but also revealed variations in breathing depth, which were not readily apparent in the visual data. In the scene, the measured respiration rate was approximately 80 breaths per minute, which is consistent with past observations (Fahlman et al., 2005).

**Fig. 4.**
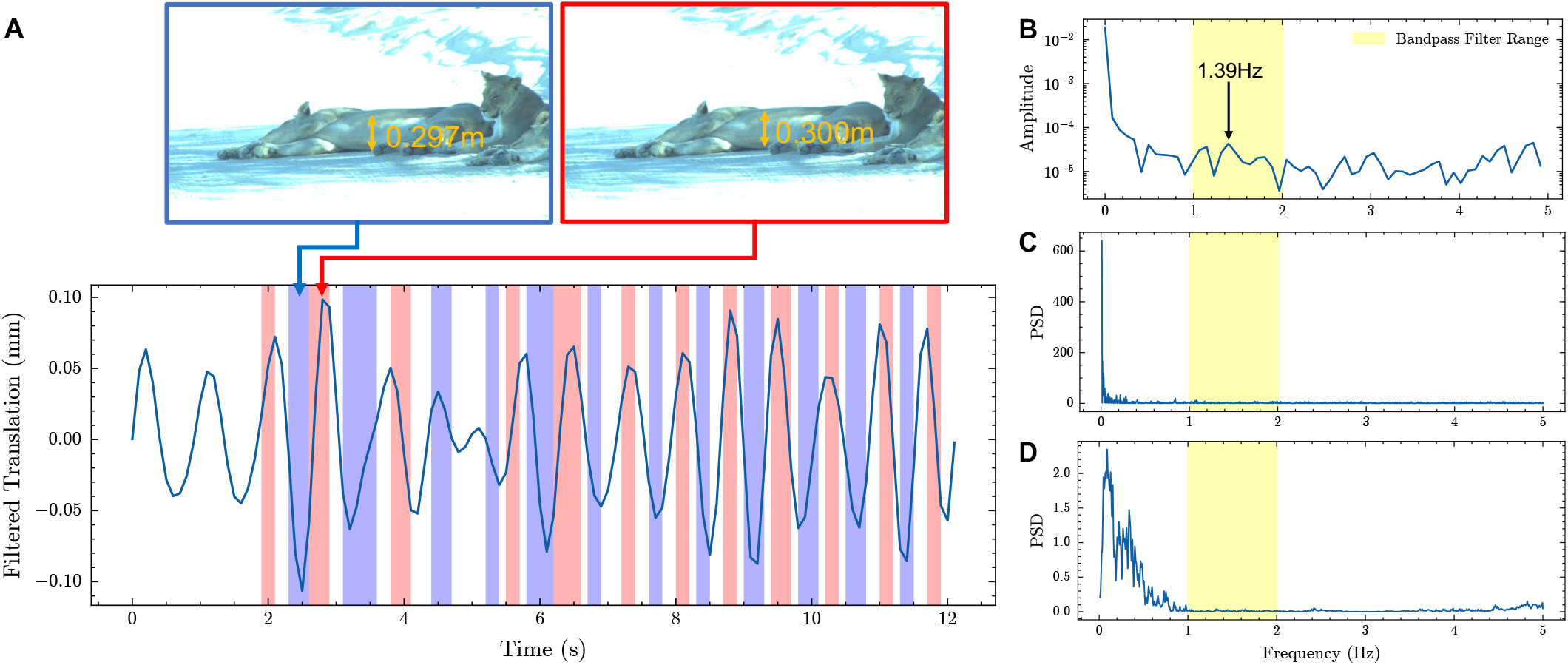
Respiration pattern analysis of a sleeping female lion using video frames and point cloud data. (A) The bottom panel shows the extracted respiratory waveform, displaying the filtered translation of the female lion’s chest over time. The y-axis represents the filtered chest movement, while the x-axis shows time. Colored shadows indicate human-labeled phases of inhalation (red) and exhalation (blue). The top panels present two sample frames of the sleeping lion at minimum and maximum filtered translation values, with corresponding chest heights from the ground obtained from the median value of the point cloud frame.(B) The frequency domain representation of the chest translation. The strongest peaks at 1.39 Hz is visible within the respiration frequency range of 1–2 Hz (yellow shaded area). (C) The frequency domain representation of the acceleration data from the IMU installed on the LiDAR, illustrating hardware vibration. (D) The frequency domain representation of the pitch velocity angle from the IMU, further characterizing system vibrations. In both (C)and (D), the yellow shaded area indicates the 1–2 Hz passband of the filter used to isolate the respiratory signal from various noise sources.

Moreover, the LiDAR-based breath measurement could capture the relative pattern transition, similar to invasive sensors (Czapan-skiy et al., 2022). An analysis of the video data suggested that the lion’s breathing was initially shallow and irregular between 4 and 5 seconds, stabilizing into a more consistent and deep breathing pattern afterward. This observation was supported by the plotted data, which also showed relatively small thoracic motion in the initial interval, becoming more pronounced and regular subsequently.

Fig. 4B shows the frequency domain of the chest translation, revealing the top peak at 1.39 Hz within the respiration frequency range (1–2 Hz according to Rzucidlo et al. (2023)). While the band-pass filter of 1–2 Hz was applied to isolate the breathing frequency, noise exists across the entire frequency spectrum. To investigate the effects of system vibration on the measurements, Fig. 4C and D present the frequency spectra of acceleration and pitch velocity data from the IMU installed in the LiDAR, revealing noise outside the respiratory band. These vibrations would be caused by various factors such as strong wind, car engine vibration, and potentially thermal effects or atmospheric disturbances. The band-pass filter successfully removed these noise sources from the respiratory signal. Higher frequency noise, particularly evident above 2 Hz, could be attributed to measurement errors, especially considering the inconsistent LiDAR point pattern across frames. Lower frequency noise (below 1 Hz) might be due to slower environmental changes or subtle movements of the lion. The amplitude of these noise sources varies, with hardware vibrations showing lower amplitudes in Fig. 4D, while measurement errors contribute smaller amplitudes across a broader spectrum (Fig. 4B).

## Conclusions

WildPose serves as a long-range 3D motion capture system, enabling researchers to study animals and their behavior within their natural habitats at an unprecedented level. Its capacity for behavioral recording through a portable system unlocks avenues for diverse experiments, from investigating individual interactions within a herd to addressing broader concerns in wildlife health management.

While WildPose has such inherent advantages, there is room for improvement in the current configuration. In its current iteration, the system relies heavily on manual controls, making data quality contingent on the operator’s expertise. In terms of data acquisition, incorporating multiple camera types could enhance the value of the data captured, as observed in previous motion capture systems designed for free movement (Nourizonoz et al., 2020). For instance, we encountered scenarios where the narrow field-of-view of a zoom-lens camera was insufficient for capturing behaviors, such as a cheetah spotting a springbok from approximately 30 meters away. The addition of a wide-view camera could mitigate such limitations.

Regarding post-processing, the system offers ample scope for improvements. The LiDAR’s laser beams are occasionally obstructed by tall grass in savanna environments, compromising 3D position accuracy near an animal’s feet. Advanced algorithms, potentially leveraging machine learning, are needed to offset the limitations inherent to each sensor. Because this challenge is difficult to address solely through post-processing —especially when grass may not appear in the video due to the zoom-lens camera’s narrow depth of field— it remains a focal point for future research.

Another significant data-processing challenge arises from the disparity between the sensors on WildPose. While the sensor frame rate is constrained by the lowest one, in our case LiDAR with 10 FPS, compared to the camera’s 170 FPS, in the current process to generate the colored point cloud, advances in multimodal sensor fusion techniques, similar to those used in autonomous driving (Almalioglu et al., 2022), may offer a potential solution.

The system’s robust design, complemented by its active gimbal and remote connectivity, positions it as a useful tool for capturing genuine behaviors in expansive and natural settings. Its portability also extends the potential for multi-view studies of wild animal behavior. Moreover, WildPose holds promise in the realm of wildlife health monitoring. Algorithms that have proven effective in livestock health monitoring through computer vision and depth cameras in indoor settings (Fernandes et al., 2020; Huang et al., 2018; Pezzuolo et al., 2019) could be adapted for use with WildPose.

In summary, WildPose represents a critical technological advancement, integrating LiDAR and zoom-lens cameras to fill the existing void between lab-based motion capture and field study instrumentation. We envision this system pioneering a new frontier in the reliable collection of wildlife motion data, thereby catalyzing future interdisciplinary research efforts.

## LIST OF SYMBOLS AND ABBREVIATIONS

LiDAR: Light Detection and Ranging
FOV: Field of View
CAD: Computer-Aided Design
ROS2: Robot Operating System 2
FPS: Frames Per Second
SAM: Segment Anything Model
RANSAC: Random Sample Consensus
IMU: Inertial Measurement Unit
PSD: Power Spectral Density
r: Radius of circular path
d: Distance between the system and tracked object
p: Subject (person) position
c: the center of circular path
X: WildPose system position

## Acknowledgements

We thank all people that helped in designing and setting up the hardware and during the fieldwork procedure, especially Nkabeng Mzileni, Correra Links and Michael Katsoulis.

## Contribution

AP and NM conceptualized the system. NM designed and developed the hardware system. NM designed and developed the general reconstruction and analysis framework. SS and NM desgined and wrote the segmentation and tracking software. NM, AP andAM designed all the case studies. NM performed the field work and the experiment and captured all the data. NM analyzed data for all results. NM and SS wrote the original draft of manuscript. NM, SS, QD, AM and AP reviewed and edited the manuscript.

## Data availability

WildPose software can be found at https://github.com/DenDen047/WildPose_v1.1/tree/main/wildpose. The analysis and application code can be found at https://github.com/DenDen047/WildPose_v1.1/tree/main/applications. All relevant data can be found within the article and its supplementary information.

