## Supplementary Information for "WildPose: A Long-Range 3D Wildlife Motion Capture System"

© 2019. MANUSCRIPT SUBMITTED TO JOURNAL OF EXPERIMENTAL BIOLOGY (2019) 00, jebxxxxxx. doi:10.1242/jeb.xxxxxx

METHODS & TECHNIQUES

### MOVIE LEGENDS

**Movie 1**. Animation of the heat map of the distances between each pair of individual springboks over time. Warmer colors indicate shorter distances and cooler colors indicate longer distances. Empty cell means that both individuals were not present at the same time simultaneously. This animation provides a dynamic visualization of the spatial relationships and social interactions within the herd.

**Movie 2**. The respiration pattern of a female lion. The left panel shows the estimated respiration pattern, which is identical to the plot in Fig. 4, while the right panel displays the raw video footage of the lion.

[1]African Robotics Unit, University of Cape Town, Cape Town, 7700, Western Cape, South Africa
[2]Department of Computer Science, University of Oxford, 7 Parks Road, Oxford, OX1 3QG, United Kingdom

Journal of Experimental Biology (2019) 00, jebxxxxxx. doi:10.1242/jeb.xxxxxx

**Table S1. Comparison of approaches for recording free-ranging animal behavior.**

| Approach | Invasiveness | Morphometrics | Portability | 3D whole-body data | Tracking volume/-duration |
|---|---|---|---|---|---|
| GPS-IMU collar (Wilson et al., 2018) | Requires animal capture and sedation to fit collar | Limited to coarse body measurements | Highly portable, suitable for long-range tracking | No 3D body pose or surface contour data | Large areas over extended periods |
| Static multi-camera field videography (Theriault et al., 2014) | Non-invasive, does not require animal manipulation | Can provide some morphometric data | Fixed installation, not easily portable | Can capture 3D data within camera overlap zone | Limited to camera field of view |
| Camera trap (Cui et al., 2020; Klasen and Steinhage, 2022) | Non-invasive monitoring | Basic morphometrics possible with reference objects | Fixed installation at specific locations | Depth-sensing traps can provide coarse 3D data in close distance | Restricted to high-traffic areas like trails or waterholes |
| UAV (Basu et al., 2019) | Non-invasive, but may alter behavior due to noise | Can estimate some morphometrics from aerial views | Highly portable, but limited flight time | No detailed 3D body data | Moderate area coverage, duration limited by battery life |
| Zoom lens photography (Brown and Wells, 2020) | Non-invasive, long-range observation | Can measure some morphometrics with scene references | Highly portable handheld or tripod setup | No 3D data, limited to 2D images | Flexible, but no path data between images |
| Telephoto lens stereo vision (de Margerie et al., 2015) | Non-invasive, long-range stereo imaging | Limited morphometrics at a distance | Portable but bulky dual-camera setup | Coarse 3D position, but no detailed surface data | $< 100\,\mathrm{m}$ range, accuracy $< 0.1\,\mathrm{m}$, no long-term tracking |
| **WildPose (ours)** | Non-invasive, long-range LiDAR scanning | Detailed 3D morphometrics of surface and pose | Portable handheld or tripod setup | Metric 3D point clouds of full body but sparsity is changed for the situation | 18–50 m range, 11–50 mm accuracy, point density increases with stationary time |

Journal of Experimental Biology (2019) 00, jebxxxxxx. doi:10.1242/jeb.xxxxxx

**Table S2.** Comparison of Depth Sensing Devices

| Device | FOV | Depth Range | Size (mm) | Frame Rate | Depth Accuracy |
|---|---|---|---|---|---|
| Intel RealSense D457 | 90° × 65° (RGB)<br>87° × 58° (Depth) | 0.6 – 6 m | 124 × 29 × 36 | 30 fps (RGB)<br>90 fps (Depth) | <2% at 4 m |
| Microsoft Azure Kinect DK | 90° × 74.3°(RGB)<br>75° × 65° (Narrow)<br>120° × 120° (Wide) | 0.5 – 5.46 m | 103 × 39 × 126 | 30 fps | <0.2% of range |
| Stereolabs ZED X | 110° × 80° × 120° (2.2 mm)<br>80° × 52° × 91° (4 mm) | 0.3 – 20 m (2.2 mm lens)<br>1 – 35 m (4 mm lens) | 164 × 32 × 37 | 120 fps | 3% at 8 m |
| WildPose | 38.8° × 21.2° – 4.03° × 2.15° (Camera)<br>14.5° × 16.2° (LiDAR) | 2 – 200 m | 290 × 290 × 530 | 480,000 points/s (LiDAR)<br>170 fps (RGB) | 2 cm at 20 m<br>Angular: < 0.03° |

**Table S3. Positioning error analysis for static individuals.** Each item indicates the measured mean and standard deviation of the positioning errors in the x, y, and z axes for two static individuals (ID 4 and ID 8).

| | ID 4 | ID 8 |
|---|---|---|
| $x$ | $-2.039 \pm 0.105\,\mathrm{m}$ | $-2.871 \pm 0.088\,\mathrm{m}$ |
| $y$ | $0.429 \pm 0.039\,\mathrm{m}$ | $0.525 \pm 0.028\,\mathrm{m}$ |
| $z$ | $140.020 \pm 0.023\,\mathrm{m}$ | $139.193 \pm 0.028\,\mathrm{m}$ |

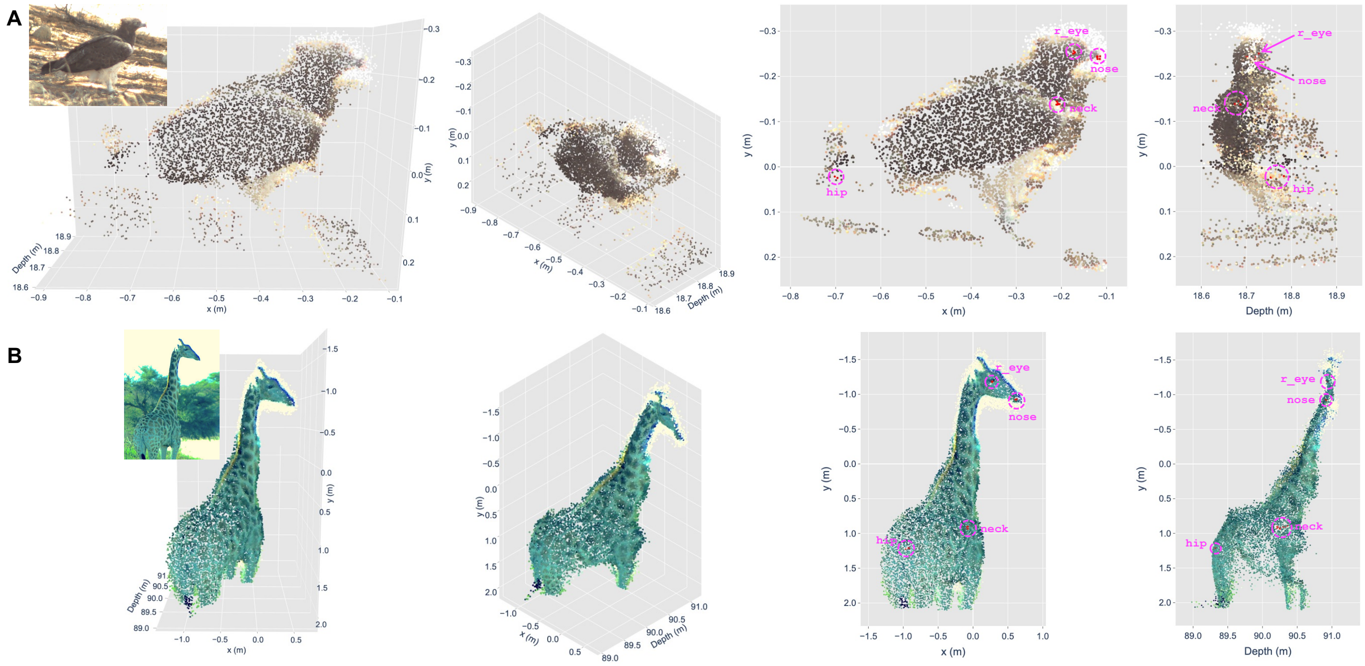


**Fig. S1. Colored point clouds of wildlife.** Dense point clouds and corresponding camera frames are shown for static animals: (A) a martial eagle and (B) a giraffe. For each scene, 20 LiDAR frames (2 seconds) were accumulated. The first column presents the colored 3D point cloud alongside the corresponding image frame, while the second column offers an oblique view of the point cloud. The third and fourth columns depict X-Y and Depth-Y plane views, respectively. Red points represent the projected 3D points of manually selected 2D keypoints in each frame, with their respective labels in pink.

Journal of Experimental Biology (2019) 00, jebxxxxxx. doi:10.1242/jeb.xxxxxx

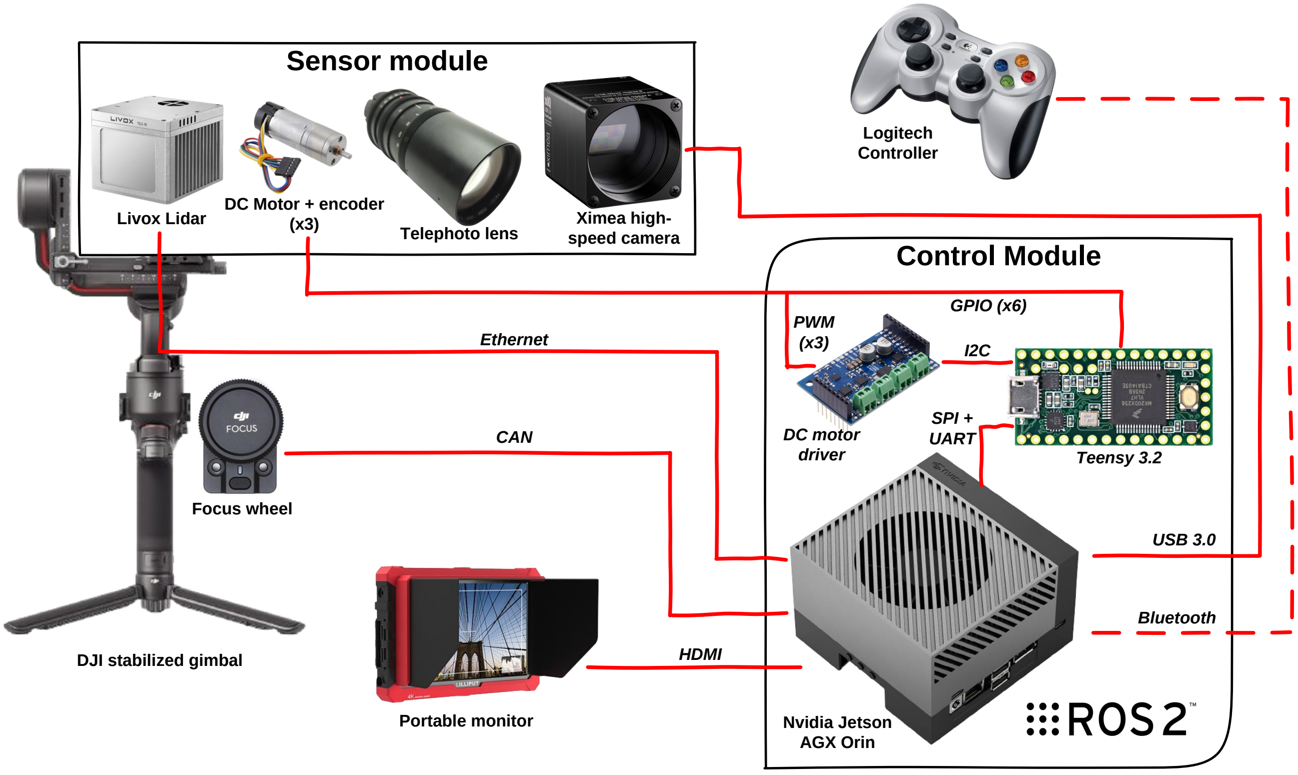


**Fig. S2. An overview of the hardware sub-systems which comprise WildPose.** The sensor module contains primary sensors, the solid-state lidar (Livox Tele-15) and the high-speed camera (Ximea MQ022CG-CM), connected to a zoom lens driven by three brushed DC motors for zoom, focus and aperture. These are housed on a stabilized gimbal (DJI RS3 Pro). The control module contains an NVIDIA Jetson AGX Orin with customized software running on the ROS 2 Foxy and handles the time-synchronization, data logging, motion control of the gimbal as well as real-time display of the video on a portable monitor. The microcontroller (Teensy 3.2) handles the low-level hardware interfacing to the motors.
